# Elevated urea levels in human frontotemporal dementia and amyotrophic lateral sclerosis *post-mortem* brain tissue: Evidence of a multi-dementia pathogenic mechanism

**DOI:** 10.1101/2025.11.03.686199

**Authors:** Sasha A Philbert, Stephanie J Church, Richard D Unwin, Garth J S Cooper

## Abstract

Frontotemporal dementia (FTD) and amyotrophic lateral sclerosis (ALS) represent two neurodegenerative diseases on opposite sides of a movement disorder continuum. However, like many other neurodegenerative diseases, the molecular pathogenesis of FTD and ALS is not fully understood. Our group has previously reported evidence for a pervasive elevation of brain urea levels in five other dementia-causing diseases. However, brain urea levels have yet to be measured in ALS and FTD. Here, we employed ultra-high-performance liquid chromatography-tandem mass spectrometry to characterise brain urea differences between control (*n*=14/12) and ALS/FTD (FTD: *n*=8/9; ALS: *n*=13/14) cases in *post-mortem* tissue from two brain regions with different levels of neuropathological burden (high vs. low). Elevated urea levels were observed in both the frontal cortex (high neuropathological burden) and primary visual cortex (low neuropathological burden) in cases with FTD. Contrastingly, in cases with ALS, elevated urea was observed in the primary motor cortex (high neuropathological burden), but not the dentate nucleus (low neuropathological burden). These results not only suggest that elevated urea levels are also present in ALS and FTD but imply that elevated brain urea is linked to a multi-dementia pathogenic mechanism. In contrast to ALS, the observation of elevated urea in regions of both high and low neuropathological burden in FTD implies that this phenotype is likely widespread and, therefore, may play a larger role in the pathogenesis of disease. Such a mechanism could offer new directions for developing treatments targeting this underlying pathology.

## 2 Introduction

Both frontotemporal dementia (FTD) and amyotrophic lateral sclerosis (ALS) are devastating neurodegenerative diseases, with estimated incidence rates of 15.1 (in the 45–65 years age group) and 2.1 new cases per 100,000 per year, respectively^1,2^. FTD is considered the second most prevalent age-related dementia in individuals under 65 and is a complex, heterogeneous disorder^3^. In FTD patients, non-Alzheimer-type neuropathological entities, collectively known as frontotemporal lobar degeneration (FTLD), vary depending on the clinical syndrome but may include degeneration of the frontal and/or anterior temporal cortices^4^. Conversely, ALS is a rare and fatal neurodegenerative disease characterised by muscle weakness due to dysfunction of the upper and lower motor neurons, primarily in the motor cortex and spinal cord^5^. Additionally, ALS patients often exhibit mild whole-brain atrophy and frontotemporal atrophy^6^.

Given the well-documented clinical overlap between ALS and FTD, both diseases are now understood to exist on opposite ends of an ALS–FTD spectrum. For example, a large-scale study found that 15% of ALS patients met the criteria for FTD, and 50% showed cognitive impairment, including executive dysfunction^7^. At the genetic level, several mutations have been linked to both diseases^8^. Similarly, molecular evidence has shown that TAR DNA-binding protein of 43 kDa (TDP-43)-positive cytoplasmic inclusions are major pathogenic hallmarks of both ALS and FTLD^9-11^. Despite recent advances in understanding these diseases, the molecular pathogenesis of ALS and FTD/FTLD remains unclear, and there are currently no effective therapeutic strategies for either condition^12,13^.

Our research group previously identified elevated brain urea levels in Huntington’s disease^14,15^, and further studies have shown the same phenotype in five other dementia-causing diseases, including Alzheimer’s disease^16^, Parkinson’s disease dementia^17^, dementia with Lewy bodies^18^, and vascular dementia^19^. Others have also shown dysregulation of urea in late-stage Huntington’s disease CSF^20^ and urea-cycle intermediates in various dementias^21-23^. These findings suggest that the presence of elevated brain urea is a novel multi-dementia phenotype and highlight urea production or clearance as a potential universal therapeutic target for dementia. However, brain urea levels have not yet been explored in ALS and FTD. In this study, we used ultra-high-performance liquid chromatography-tandem mass spectrometry (UHPLC-MS/MS) to assess case–control differences in post-mortem brain urea levels across regions with varying neuropathological burdens in ALS and FTD. Elevated brain urea levels were observed in both FTD and ALS. While this urea phenotype appears to be widespread in FTD (affecting both high- and low-burden brain regions), it seems more localised in ALS, primarily in the high-severity region, aligning with the disease’s progression. Overall, these results provide evidence for novel patterns of brain urea perturbations in FTD and ALS, highlighting the role of a urea-mediated mechanism in now seven age-related dementias.

## 3 Materials and Methods

### 3.1 Case selection

To conduct a case–control investigation of ALS and FTD, human *post-mortem* brain tissues were obtained from the University of Maryland Brain and Tissue Bank and the Human Brain and Spinal Fluid Resource Centre (part of the NIH NeuroBioBank), respectively. For the ALS case–control investigation, 14 controls and 14 ALS cases were sampled. For the FTD case–control investigation, 14 controls and 9/8 FTD cases were sampled. One FTD case in the frontal cortex only was unable to be analysed due to an insufficient amount of tissue. All cohorts were matched for age, sex, and *post-mortem* interval (PMI). The inclusion/exclusion criteria for both FTD and ALS cases included a confirmed neuropathological diagnosis of FTD/ALS, PMI < 30 h, and no histopathological findings of other diseases likely to cause dementia (e.g., Alzheimer’s disease). Controls had no history of dementia and no other neurological abnormalities. Mean PMI was ≤ 24 h for ALS/FTD cases and controls (**Tables 1 & 2**); importantly, extended *post-mortem* delay of up to 72 h has been shown not to interfere with brain-urea levels^24^.

**Table 1.** Group characteristics for the FTD cohort.

| Variable | Control | FTD (PVC) | FTD (FC) |
| --- | --- | --- | --- |
| Number | 14 | 9 | 8 |
| Age | 70 (68–73) | 70 (63–78) | 68 (61–76) |
| Male sex, *n* (%) | 3 (21) | 2 (22) | 2 (25) |
| Post-mortem interval (h) | 22 (20–24) | 23 (20–26) | 24 (20–27) |

Values are mean ( $\pm 95\%$ CI) age and post-mortem intervals for each group. Exact male/female ratios could not be maintained across all groups due to differences in group numbers. All other cohort differences were non-significant.

For this initial study, sets of tissue from two brain regions per disease were chosen, each with different levels of neuropathological burden (low vs. high). For both diseases, the degree of neuropathological burden was based on the severity of TDP-43 pathology and brain atrophy. For ALS, the primary motor cortex (Brodmann area 4; high neuropathological burden) and the dentate gyrus (low neuropathological burden) were chosen. For FTD, the inferior frontal cortex (Brodmann area 11; high neuropathological burden) and the primary visual cortex (Brodmann area 17; low neuropathological burden) were chosen. The sampling of these regions was also chosen to enable comparisons with brain urea measurements from our group’s previous dementia datasets.

### 3.2 Tissue dissection

All samples were transported from the respective NIH NeuroBioBank repositories to the University of Manchester on dry ice and then stored at −80°C. Samples were thawed briefly on ice before being cut into 20 mg aliquots (±2 mg; wet weight) using a ceramic scalpel and placed into 2 mL “Safe-Lock” microcentrifuge Eppendorf tubes (Eppendorf AG; Hamburg, Germany). Samples were stored at −80°C pending metabolite extraction.

### 3.3 Metabolite extraction

All brain samples were extracted in 800 μL 50:50 (v/v) methanol:chloroform mixture, which contained 1mM stable isotopically-labelled urea internal standard (Urea-^15^N_2_, Sigma–Aldrich) and had been maintained at −20°C for a minimum of 4 h. Tissue samples were placed in a TissueLyser plate (stored at −80°C for an hour before use) and extracted at 25 Hz for 10 min with a single 3 mm-tungsten carbide bead per sample in a TissueLyser bead homogeniser (Qiagen, Manchester, UK). After extraction, 400 µL of LC-MS grade water was added to each sample before it was briefly vortexed and then centrifuged at 2,400 × g for 10 min to ensure phase separation. A total of 200 μL of the polar methanol phase from each sample was then transferred to a new pre-labelled microcentrifuge tube and dried overnight using a centrifugal concentrator (Savant SpeedVac™; Thermo-Fisher, Waltham, MA). Once dried, 400 μL of 0.1% formic acid was added to each sample before it was briefly vortexed. Then, 100 μL of the resulting solution was transferred to 300 μL-insert autosampler vials, along with two extraction blanks containing 100 μL 0.1% v/v formic acid only. To generate the standard curve, 10 μL of the 1mM labelled urea internal standard together with 20 uL of a 50 mM unlabelled external urea standard (urea analytical standard, 56180 Supelco, PA, United States) stock were added to 0.1% formic acid to achieve a final volume of 200 μL per standard, containing concentrations of 0–5,000 μM unlabelled urea in 0.1% v/v formic acid. Three quality control (QC) samples were also prepared containing 500 μM labelled urea and 20, 200, and 2,000 μM of unlabelled urea in 0.1% v/v formic acid.

### 3.4 Ultra-high-performance liquid chromatography-tandem mass spectrometry

Cerebral *post-mortem* urea levels were quantified via ultra-high-performance liquid chromatography-tandem mass spectrometry (UHPLC-MS/MS) using a TSQ Vantage triple quadrupole mass spectrometer coupled with an Accela UHPLC system (Thermo Fisher Scientific, MA, United States). Separation was carried out on a Hypersil Gold AQ column with a diameter of 2.1 mm, length of 100 mm, and particle size of 1.9 mm (ThermoFisher Scientific) maintained at 25°C with a 0.5 mm pre-column filter (Thermo Fisher Scientific). Gradient elution was performed using 0.1% formic acid in water (A) and 0.1% formic acid in acetonitrile (B) at 300 mL/min. At the start of the analysis, the composition was 100:0 (A:B). At 0.8 min, a 1.2 min gradient from 100:0 to 95:5 was used, followed by a 2-min gradient from 95:5 to 0:100 at 2 min. Then, an isocratic elution of 0:100 was maintained for 2 min before changing to 100:0 for 3 min. The total MS run time was 9 min per sample. Urea (parent mass = 61.1 Da; product mass = 44.1 Da) and labelled urea (parent mass = 63.1 Da; product mass = 45.1 Da) internal standards were detected using electrospray ionisation in positive ionisation mode. (**Suppl. Table 1**; **Suppl. Fig 1**)

### 3.5 Data analysis

UHPLC-MS/MS data were analysed using LCQuan software (Thermo Fisher Scientific, MA, United States). Chromatographic peaks were identified based on expected retention time (RT) and compared against labelled urea internal standard peak RTs for each QC/standard/sample. Each peak was manually checked for correct identification. Standards and QC samples were excluded from analysis if the percentage difference from the calibration curve was >15% (or 20% for the lowest standard or QC sample). At least two out of the three QC samples for each brain region had a percentage difference below this threshold. Quantification of urea in samples was performed using the ratio of urea peak area to internal standard peak area and comparison to the standard curve. Brain-urea concentrations were first exported to Excel and then corrected for sample weight and dilution, and finally converted to units of mmol/kg. The significance of brain urea case–control differences was determined using either Welch’s t-tests or Mann–Whitney U tests depending on whether or not data were found to be normally distributed using the Shapiro–Wilks test of normality. FTD/ALS case-variable differences were determined using the Brown–Forsythe and Welch analysis of variance (ANOVA) test with Dunnett’s T3 multiple comparisons. Statistical calculations were performed using GraphPad Prism v9.2.0 (GraphPad; La Jolla, CA). All tests were two-sided. *p*-values<0.05 were considered significant and *p*-values <0.10 have also been tabulated. Post-hoc statistical power and sample-size estimates were calculated using G*Power v3.1.9.4^25^.

### 3.6 Sensitivity analysis

Sensitivity analyses were employed to further understand the strength of association to potential or uncontrolled confounding variables. Box plots were fitted using SPSS (version 29.0.0.0) to identify outliers in all urea datasets (**Suppl. Fig. 2**), and the impact of distributional assumptions was determined by comparing data from both parametric and non-parametric methods^26^.

The risk ratio (RR), also known as relative risk, was calculated using the following formula:

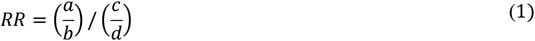

where **a** and **c** correspond to the number of cases and controls with urea levels >95% of the upper confidence interval (CI) limit of the controls, and **b** and **d** correspond to the total number of cases and controls in the cohort, respectively.

The E-value can be interpreted as “the minimum strength of association, on the risk ratio scale, that an unmeasured confounder would need to have with both the treatment and the outcome to fully explain away a specific treatment-outcome association, conditional on the measured covariates”^27^. E-values were calculated using the following formula:

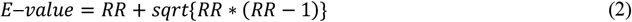

The following formula was used to calculate the E-value of the RR CIs:

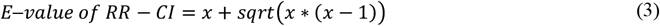

where **x** is the CI closest to 1. If the confidence intervals crossed 1, the E-value of the RR-CI was recorded as 1.

Although the *p*-value is necessary to confirm whether an effect is present, the effect size is key to understanding the magnitude of the difference between groups. Here, Glass’s delta was applied, rather than Cohen’s *d*, due to differences in standard deviations between cases and controls. Glass’s delta was calculated using the following formula:

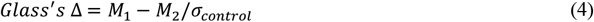

where **M1** is the mean case value, **M2** is the mean control value, and **σ**_**control**_ is the standard deviation of the control group. Effect size values between 0.2 and 0.5 were considered small, values between 0.5 and 0.8 were considered medium size, values between 0.8 and 1.3 were considered large, and values >1.3 were considered very large.

As *p-*values on their own can be subject to misinterpretation, S-values (or Shannon information values) have been calculated to provide a preferable scale of information in ‘bits’ against the experimental hypothesis^28^. S-values were calculated using the following formula:

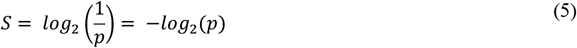

## 4 Results

### 4.1 Case characteristics

Tissues were obtained from two brain regions (one high-severity and one low-severity region) in ALS (*n*=14/13) and FTD (*n*=8/9) cases. Corresponding controls with similar ages and PMIs were sampled from both ALS (*n*=12) and FTD (*n*=14) case–control investigations. Although one control (BBN: S14842) received a diagnosis of ‘unaffected control’, the neuropathological assessment was consistent with early-stage vascular dementia (**Suppl. Table 2**). Therefore, this control was excluded from this study as our group’s previous evidence suggests that vascular dementia can independently cause elevated brain urea levels^19^. Following the fitting of box plots for sensitivity analysis, two controls (BBN: 6688; BBN: 6819) and one ALS case (BBN: 5789) were identified as extreme outliers and excluded from the study (**Suppl. Table 3 & Suppl. Fig. 2**). Extreme outliers were defined as being more than 3 interquartile range (IQR) below the first quartile or above the third quartile. No extreme outliers were identified in the FTD case–control investigation and mild outliers (1.5 IQR below the first quartile or above the third quartile) have been included in the following analyses.

One ALS case (BBN: 6350) was diagnosed with familial ALS (*C9orf72*-positive). Although most ALS cases are sporadic, ∼10–15% of ALS cases are familial^28^. However, despite the differences between familial and sporadic ALS, this case did not present any outliers in either brain region. Likewise, one FTD case (BBN: S18571) was diagnosed with valosin-containing protein disease caused by mutations in the *VCP* gene. Neuropathologic evaluation of this case revealed a TDP-43 proteinopathy consistent with FTLD-TDP-43. No other FTD or ALS cases had genetic diagnoses. Although efforts were made to ensure the consistency of cohort variables across all groups, after removing outliers, the mean age for the FTD controls (70 y [95% CI=68–73]) was lower than the ALS controls (54 y [95% CI=47–72]; p = 0.03; **Suppl. Fig. 3**). However, no differences in urea were observed between these two control cohorts (**Suppl. Fig. 4**). When separated by gender, female ALS cases were older than male controls (**Suppl. Fig. 5**). No other significant case–control or inter-cohort differences for *post-mortem* interval (PMI) or age were present in these tissues (**Tables 1 & 2; Suppl. Fig. 3**).

**Table 2.** Group characteristics for the ALS cohort.

| Variable | Control | ALS (DN) | ALS (PMC) |
| --- | --- | --- | --- |
| Number | 12 | 14 | 13 |
| Age | 54 (47–72) | 67 (47–72) | 67 (59–75) |
| Male sex, *n* (%) | 8 (67) | 9 (64) | 9 (69) |
| Post-mortem interval (h) | 21 (18–24) | 19 (19–23) | 19 (15–23) |

Values are mean ( $\pm 95\%$ CI) age and post-mortem intervals for each group. Exact male/female ratios could not be maintained across all groups due to differences in group numbers. All other cohort differences were non-significant.

### 4.2 Brain urea levels

Statistically significant urea elevations were observed using UHPLC-MS/MS in FTD cases for both high- and low-severity brain regions, but only the high-severity region in ALS cases. In the FTD case–control investigation, FTD cases displayed mean fold increases of 2.39 and 2.08 in the frontal cortex (high-severity region; p=0.005) and primary visual cortex (low-severity regions; p=0.007), respectively (**Table 3; Fig. 1**). In the ALS case–control investigation, ALS cases displayed a fold change of 1.75 in the primary motor cortex (high-severity region; p=0.016). Despite showing a 1.99-fold change in the mean, differences between ALS cases and controls in the dentate nucleus (low-severity region) were not statistically significant (p=0.212; **Table 4; Fig. 1**). It is worth noting that one control sample in the dentate nucleus was considerably higher than the others, which may have skewed the data. Interestingly, mean urea levels were approximately 2-fold higher in the primary motor cortex than in the dentate nucleus for controls and ALS cases.

**Fig 1.**
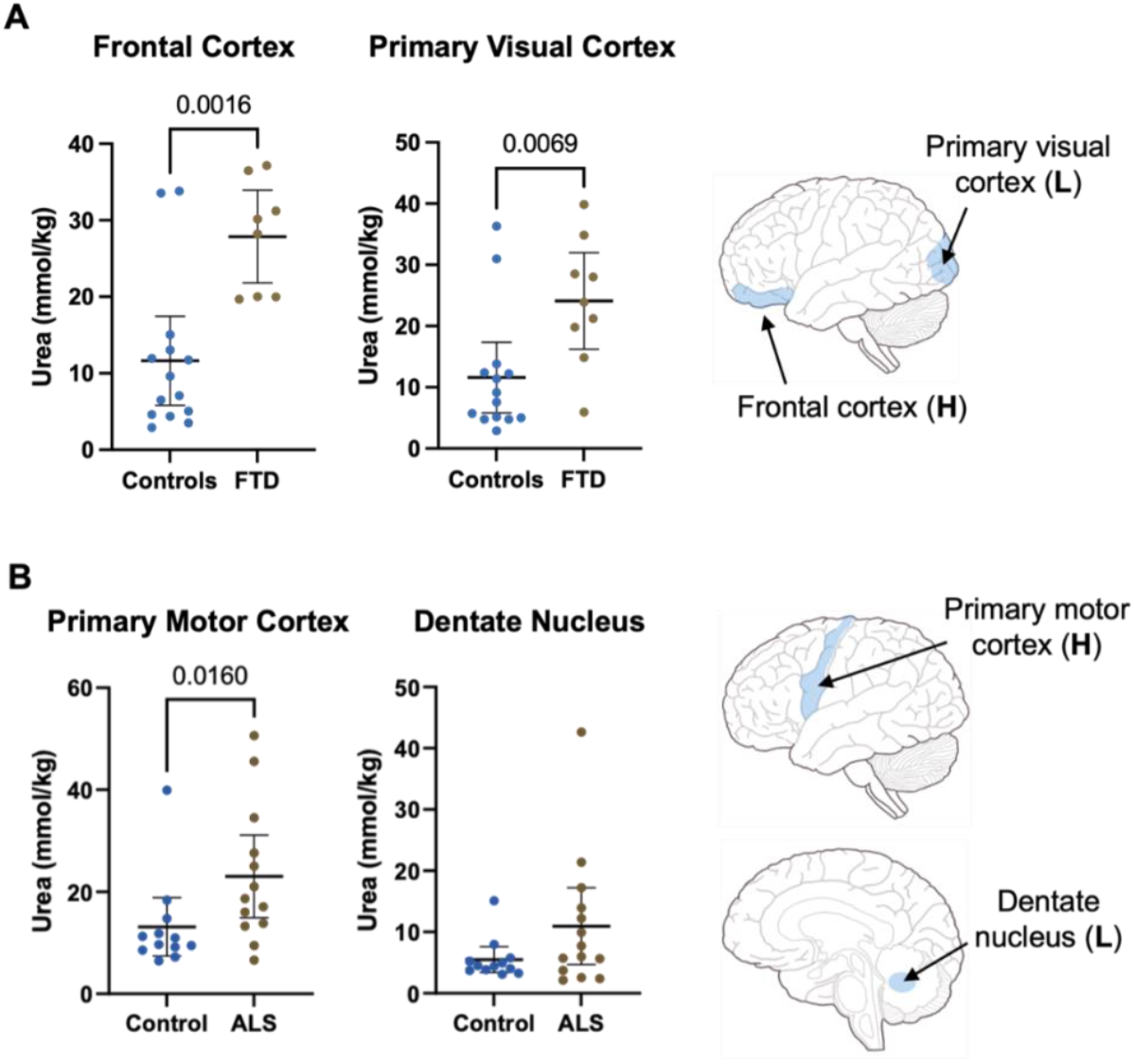
Regional brain urea concentrations for **A)** FTD case–control and **B)** ALS case–control *post-mortem* tissue comparisons. Data are means ±95% CI. *p*-values for the significance of individual region between-group differences were calculated using Mann–Whitney U tests based on urea measurements from FTD case–control (con=15 vs. FTD=8/9) and ALS case–control (con=14 vs. ALS=14) investigations. Abbreviations: H: High neuropathological burden; L: Low neuropathological burden.

**Table 3.** Urea concentrations for controls and FTD cases.

| Brain region | Control (mmol/kg) | FTD (mmol/kg) | Fold-change | Welch's t-test | Mann–Whitney U test |
| --- | --- | --- | --- | --- | --- |
| Frontal cortex | 11.64 (5.80–17.46) | 27.86 (21.81–33.91) | 2.39 | 0.0003 | 0.0016 |
| Primary visual cortex | 11.59 (5.81–17.37) | 24.11 (16.23–32.00) | 2.08 | 0.010 | 0.007 |

Data are means ( $\pm 95\%$ CI). *p*-values for the significance of individual region between-group differences were calculated by Welch's t-test and Mann–Whitney U test based on urea measurements from controls (*n*=15) and FTD (*n*=8/9) brains.

**Table 4.** Urea concentrations for controls and ALS cases.

| **Brain region** | **Control**<br>**(mmol/kg)** | **ALS**<br>**(mmol/kg)** | **Fold-**<br>**change** | **Welch's**<br>**t-test** | **Mann–Whitney**<br>**U test** |
| --- | --- | --- | --- | --- | --- |
| Primary motor cortex | 13.15 (7.40–18.89) | 23.02 (14.90–31.15) | 1.75 | 0.042 | 0.016 |
| Dentate nucleus | 5.50 (3.40–7.59) | 10.95 (4.69–17.21) | 1.99 | 0.093 | 0.212 |

Data are means ( $\pm 95\%$ CI). *p*-values for the significance of individual region between-group differences were calculated by Welch's t-test and Mann–Whitney U test based on urea measurements from controls (*n*=14) and ALS (*n*=14) brains.

When separated by sex, urea levels in the female control group (10.75 mmol/kg) were lower than in the female FTD group (28.8 mmol/kg; p=0.024) in the frontal cortex. Urea levels in the male control group (10.25 mmol/kg) were also lower than in the male ALS group (26.32 mmol/kg; p=0.012) in the primary motor cortex (**Suppl. Fig. 6**). No other sex differences were observed in the remaining groups or brain regions. However, it should be noted that the male–female ratio for the ALS case–control comparison was heavily skewed towards females (males=5 subjects; females=18 subjects).

*Post-hoc* sample size estimates were computed to further ensure the reliability of these case–control differences. For FTD, only the frontal cortex met the required sample size (*n*=18). Following the removal of outliers, the primary motor cortex was just below the required sample size (*n*=24). For ALS, the estimated sample sizes for the primary motor cortex and dentate nucleus were *n*=72 and *n*=44, respectively.

### 4.3 Sensitivity analysis

The fitting of box plots confirmed the presence of two control outliers in the primary motor cortex and dentate nucleus control and one ALS case outlier in the primary motor cortex only. These outliers were removed before the assessment of urea level differences. No outliers were observed in the FTD case–control investigation for both regions. To assess the robustness of these data under different distributional assumptions, both parametric (Welch’s t-test) and non-parametric (Mann–Whitney U test) analyses were conducted. Three regions (frontal cortex, primary visual cortex, and primary motor cortex) presented statistically significant urea differences using both Welch’s t-tests and Mann–Whitney U tests.

For FTD case–control comparisons, the RR and E-values were 6.13 and 11.7, respectively, for the frontal cortex, and 5.0 and 9.5, respectively, for the primary motor cortex. Effect sizes, measured by Glass’s Δ, were 1.3 for both regions, which are considered large. Parametric and non-parametric S-values in the frontal cortex were 8.2 and 7.6, respectively, and 6.6 and 7.2, respectively, in the primary visual cortex. For ALS case-control comparisons, the RR and E-values were 5.54 and 10.6, respectively, for the primary motor cortex, and 3.00 and 5.4, respectively, for the dentate gyrus. The Primary motor cortex displayed an effect size of 1.1 (large), whereas the dentated nucleus displayed an effect size of 1.7 (very large). Parametric and non-parametric S-values in the primary motor cortex were 4.6 and 6.0, respectively, and 3.4 and 2.2, respectively, in the dentate nucleus (**Suppl. Table 4**).

## 5 Discussion

Our group has previously provided evidence for a pervasive elevation of brain urea levels in five dementias. This includes Alzheimer’s disease^29^, Parkinson’s disease dementia^17^, dementia with Lewy bodies^18^, Huntington’s disease^14^, and vascular dementia^19^. However, this multi-dementia phenotype has yet to be assessed in FTD or ALS brain tissue, which represents the remaining age-related dementias. Using the same UHPLC-MS/MS methods, urea concentrations were measured in case–control comparisons of FTD and ALS. Two brain regions were sampled per comparison, reflecting either a high or low neuropathological burden. In the FTD case–control comparison, elevated brain urea concentrations were observed in both the frontal cortex (high neuropathological burden) and primary visual cortex (low neuropathological burden). However, in the ALS case–control comparison, increased urea concentrations were observed in the primary motor cortex (high neuropathological burden) but not the dentate nucleus (low neuropathological burden). No differences were observed for age or *post-mortem* intervals (PMIs), suggesting that case–control differences are unlikely to be due to these cohort variables. The lack of cohort variable effects was also observed in our group’s past dementia urea investigations for age; however, case–control differences for PMI were apparent for Parkinson’s disease dementia^30^ and Huntington’s disease^31^.

Our previous investigations of dementia-causing disease generally had cohort sizes of *n*=∼20 (10 cases vs. 10 controls); however, post-hoc power analyses revealed that this cohort size may not be sufficient for other omic analyses (e.g., trace-metal analysis^30,32,33^). Therefore, for these investigations, we aimed to obtain a sample size of *n*=28 (14 cases vs. 14 controls) to compensate for the lower power observed in earlier investigations. While this cohort size was attainable for the study of ALS, we were unable to find sufficient available FTD cases to match our inclusion/exclusion criterion (actual numbers were 8/9 FTD cases vs. 14 controls). However, this cohort size is still larger than many of our previous analyses and in the event allowed sufficient case–control comparisons of tissue urea levels.

This pattern of urea elevation in ALS is in keeping with the well-documented selective motor neuron vulnerability in the primary motor cortex of ALS patients^34,35^ and the staging of phosphorylated TDP-43 in disease^36^. The absence of urea elevation in the dentate nucleus (low pathological burden) may imply that this urea phenotype is localised to severely affected regions, such as the primary motor cortex and brain stem, even at end-stage disease. Various reasons have been proposed for this selective vulnerability of motor neurons, with calcium toxicity/excitotoxicity being the most prominent^34,37-39^. However, the precise mechanism(s) of neuronal vulnerability and regional susceptibility in ALS are still not fully understood. Urea elevations in FTD were observed in both the frontal cortex (high neuropathological burden) and primary visual cortex (low neuropathological burden), suggesting a more global brain urea phenotype by comparison, similar to our group’s previous findings in other dementias (**Fig. 2**). Therefore, a more extensive multiregional assessment of *post-mortem* brain tissue will be necessary to confirm the pattern of cerebral urea metabolism in FTD. Considering the present observations of elevated brain urea in FTD and ALS, we now have evidence of this urea phenotype in 16 out of the 18 brain regions across 7 separate age-related dementias measured by our group. The uniformity of these urea levels suggests that elevated brain urea is a multi-dementia phenotype and implies a common urea-mediated mechanism amongst the age-related dementias (**Fig. 2**). Taken together, these data highlight urea as a potential universal therapeutic target for dementia.

**Fig 2.**
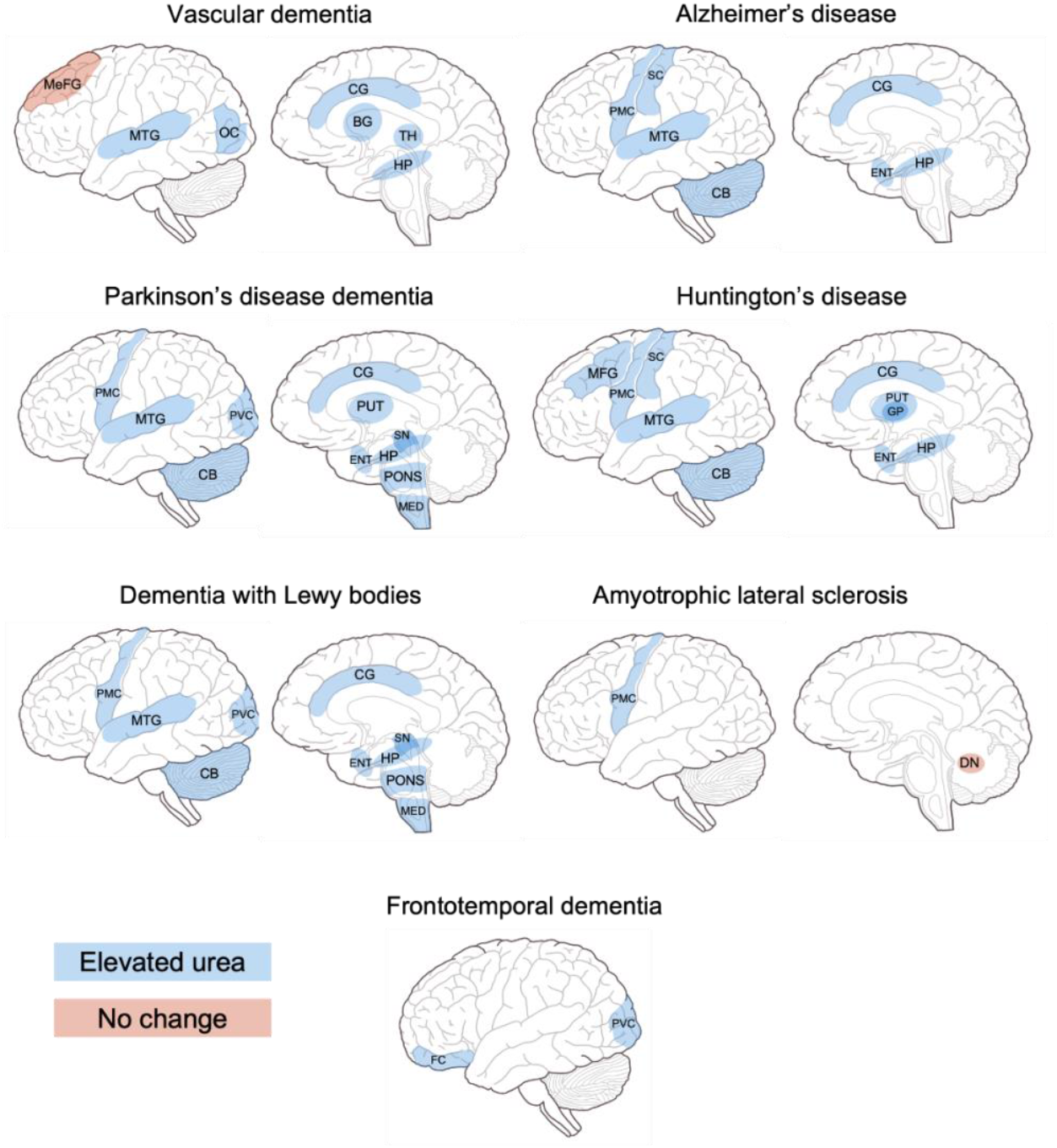
A schematic showing regional observations of brain urea in seven dementias measured by our group. Regions highlighted in blue represent increased urea levels compared to controls. Regions highlighted in orange represent no differences in urea levels. Abbreviations: BG: Basal ganglia; CB: Cerebellum; CG: Cingulate gyrus; DN: Dentate nucleus; ENT: Entorhinal cortex; FC: Frontal cortex; HP: Hippocampus; MED: Medulla oblongata; MeFG: Medial frontal gyrus; MFG: Middle frontal gyrus; MTG: Middle temporal gyrus; OC: Occipital cortex; PUT: Putamen; PMC: Primary motor cortex; PVC: Primary visual cortex; SN: Substantia nigra; TH: Thalamus.

The urea cycle functions to convert ammonia (NH_3_), a nitrogenous waste product, to urea following protein catabolism. Five enzymatic reactions are involved in the classical pathways of urea metabolism, starting with the oxidative deamination of glutamate to form ammonium ion (NH_4_^+^) and α-ketoglutarate. After synthesis, urea diffuses into the bloodstream from the liver, where it is filtered and excreted in the urine via the kidneys. When renal clearance is impaired, as seen in conditions such as chronic renal failure, urea and other uraemic toxins can build up. If left untreated, elevated systemic urea levels (i.e., uraemia) can cause uraemic encephalopathy, which can lead to dementia-like symptoms (e.g., memory loss and depression); seizures; and, in more severe cases, comas^40^. The present study shows elevated urea levels in FTD (frontal cortex = 2.39-fold-change; primary visual cortex = 2.08-fold-change) and ALS (primary motor cortex = 1.75-fold-change) *post-mortem* brain tissue. Despite the lack of research surrounding urea metabolism in these diseases, altered plasma levels of urea cycle-related analytes, homoarginine and hydroxy-proline, have been reported in human FTD and ALS studies^41,42^. However, to our knowledge, this is the first study to directly demonstrate elevated brain urea levels in FTD and ALS, suggesting a possible urea-mediated mechanism in these diseases and offering further evidence for elevated urea as a multi-dementia phenotype.

A case series examining three patients with diagnosed uremic encephalopathy revealed blood urea concentrations (converted from blood urea nitrogen values) ranging from 13.82 to 19.34 mmol/L^43^. Although measured in the periphery, these values are comparable to the brain urea concentrations presented here in FTD and ALS cases. This suggests the mechanism(s) responsible for CNS dysfunction in uremic encephalopathy may also be present in these diseases. Congenital defects of enzymes in the urea cycle, termed urea cycle disorders, can cause hyperammonaemia and subsequent brain damage^44^. Interestingly, urea cycle disorder mutations have recently been identified in non-Alzheimer’s disease and non-FTD dementia^45^, suggesting that alterations in urea cycle enzymes are directly capable of causing CNS dysfunction/dementia. Despite this, proteomic profiling of Alzheimer’s disease, which presents a similar, albeit more severe brain urea phenotype^16^, failed to identify any enrichment of urea cycle pathways^46-49^. This implies that elevated brain urea levels in dementia may not be caused by alterations in urea cycle enzymes. However, it should be noted that the role of urea cycle enzymes in FTD and ALS cannot be fully elucidated due to the current lack of corresponding proteomic investigations.

Another possible explanation for the elevated brain urea seen here is the movement of urea across the blood-brain barrier (BBB) into the CNS. Although the cationic configuration of ammonium ion prevents it from entering the CNS due to the blood–brain pH gradient, a small fraction of ammonia can readily diffuse across the BBB^50^. However, the movement of ammonia alone is insufficient to cause the elevated urea levels shown here. While urea is slow to cross the BBB owing to its hydrophilic nature, disruption to the BBB in FTD and ALS may permit urea entry into the brain^51-54^. However, neither hyperammonaemia nor uraemia, which are routinely tested for in the clinic, were reported in the metadata for any of the patients used in this study. This suggests that elevated urea probably originates in the CNS in these forms of age-related dementia.

Our group has previously suggested that increased protein catabolism, due to underlying neurodegenerative pathology, could lead to elevated brain urea levels in age-related dementias. Thus, overactivation of the urea cycle could be the primary cause of this urea phenotype. While plausible, evidence is mixed on whether a fully functional urea cycle exists in the brain. A prodromal sheep model of Huntington’s disease, which also displayed elevated brain urea levels, showed no differences or negligible expression of transcripts encoding urea cycle enzymes^31^. Likewise, elevated urea and fumaric acid, whose production is catalysed by argininosuccinase (EC 4.3.2.1), but decreased ornithine only was observed in Huntington’s disease human *post-mortem* brain tissue^14^. Although these studies suggest a partial/unaffected urea cycle in the brain, Ju et al.^22^ provide evidence for a complete urea cycle in reactive astrocytes in Alzheimer’s disease, which reverts to non-cyclic urea metabolism under normal conditions. They also identified that autophagy-mediated degradation of amyloid beta causes an accumulation of ammonia which re-enters the urea cycle, leading to elevated urea. Despite mixed evidence for the role of the urea cycle in the brain, a dichotomous cycle for FTD/ALS, similar to that shown by Ju et al.,^22^ may be present. However, more research regarding urea cycle intermediates and enzymes is required to fully test this hypothesis.

Another route for ammonia clearance in the brain is the glutamate–glutamine cycle. Predominantly occurring in astrocytes, this pathway facilitates the removal of excess ammonia via the amidation of glutamate and ammonia to glutamine^55^. Notably, human FTD iPSC-derived astrocytes have shown elevated glutamine and glutamine synthase^56^. Moreover, in ALS, increased levels of glutamate and glutamine were observed in the brain stem using proton magnetic resonance spectroscopic imaging^57^. These reports are consistent with the increased brain urea levels observed in FTD and ALS in this study, suggesting that protein catabolism may be driving this urea phenotype via the overactivation of the glutamate–glutamine cycle in response to excess ammonia. However, the question of why excess urea is not being cleared out from the CNS remains to be answered. The two main urea transporters are UT-A (*Slc14A2*) and UT-B (*Slc14A1*), of which the latter is the most dominantly expressed in the CNS^58^. Although one could assume that deficient UT-B activity might be responsible for the sustained elevation of brain urea levels in FTD and ALS, evidence from our group’s previous investigation of dementia would suggest otherwise. In human Huntington’s disease and Alzheimer’s disease *post-mortem* brain tissue, both urea concentrations and UT-B expression were higher than in controls^31,49^. As previously suggested by Philbert et al.^19^, this implies that despite increased UT-B expression, this transporter may be overwhelmed by the surplus of urea, causing a sustained pool of urea in the brain.

Carbamoylation, distinct from carbamylation, refers to the non-enzymatic post-translational modification of a primary amine or free sulfhydryl group of a protein and free amino acids with isocyanic acid to form a carbamoyl moiety (-CONH_2_)^59^. This process has been associated with increased oxidative stress, inflammation, and pro-apoptotic signalling in uraemic conditions^60-62^. Elevated urea may also lead to endothelial dysfunction via the carbamoylation of low-density lipoprotein and subsequent vascular smooth muscle cell proliferation^63^. Under normal physiological conditions, the spontaneous dissociation of urea to form cyanate and its tautomer, isocyanate, occurs at a ratio of 100:1^64^. However, during periods of uraemia, levels of cyanate and isocyanate are drastically increased. For instance, in chronic renal failure patients, plasma cyanate levels were reported to be 141 nmol/L before haemodialysis and 45 nmol/L after^65^. These findings suggest that the levels of brain urea observed here in FTD and ALS, and in other dementias investigated by our group, could well be capable of causing substantive carbamoylation and cellular dysfunction in the CNS. The present study measured *post-mortem* brain urea levels and, therefore, represents urea metabolism at end-stage disease only. Nonetheless, evidence of elevated brain urea in the prodromal phase of dementia implies that protein carbamoylation could contribute to the pathogenesis of FTD and ALS rather than simply a product of early pathogenic events^31^. If this is accurate, CNS carbamoylation might serve as a potential biomarker for early-stage dementia. However, further investigation into pre-clinical protein carbamoylation in FTD and ALS is necessary to substantiate this hypothesis.

Research on protein carbamoylation in dementia is sparse *inter alia*, partly due to challenges associated with liquid chromatography-mass spectrometry. Urea is commonly used as a denaturant in liquid chromatography-based proteomics, potentially causing additional protein carbamoylation in samples. Additionally, the mass shift of carbamoylation (+43 Da) closely resembles that of trimethylation and acetylation (+42 Da), complicating the differentiation of these modifications^66^. Magnetic resonance spectroscopy, which measures nuclear spin to provide a snapshot of the chemical environment, could be a promising alternative^67^. Magnetic resonance spectroscopy is non-invasive and does not require protein denaturation, potentially overcoming some limitations of liquid chromatography-mass spectrometry. Future studies should explore the viability of magnetic resonance spectroscopy for measuring carbamoylated lysine side chains in the brain to measure the extent of dementia neuropathology.

Sensitivity analysis was conducted to avoid misinterpretations of *p*-values and to assess the influence of potential or uncontrolled confounding variables on study results (**Suppl. Table 4**). Boxplots were initially fitted to identify and exclude potential outliers. The frontal cortex, primary visual cortex, and primary motor cortex presented statistically significant (*p* < 0.05) differences in urea using both parametric and non-parametric methods, underscoring the robustness of these associations regardless of distributional assumptions. Both regions for the investigation of FTD demonstrated relatively high s-values, reinforcing evidence against the null hypothesis. For example, the frontal cortex displayed an S-value of 12 (to the nearest integer) and, thus, can be interpreted as tossing a coin and getting the same result 12 times in a row^68^. In contrast, the s-values for ALS were 6 for the primary motor cortex and 2 for the dentate nucleus. This further illustrates that ALS likely presents no brain urea differences in the dentate nucleus at end-stage disease. The analysis of RRs and E-values for all brain regions presented ranges from 3.00–7.00 and 5.4–13.5, respectively, implying that it is unlikely that possible unmeasured confounding variables have influenced the study results.

Although this urea phenotype is regarded as a multi-dementia phenotype, one weakness of the present study is the relatively low sample size. Efforts to increase the study sample size were hampered by the exclusion of outliers and the limited tissue availability, with only the frontal cortex meeting the required *post-hoc* sample size estimate (**Suppl. Table 5**). One potential solution would be to source human tissue from multiple repositories; however, this could introduce additional bias due to variability among brain banks^69^. Another weakness is the sampling of both left and right hemispheres in the present study of FTD. Unlike ALS, FTD can present with left- or right-dominant temporal lobe atrophy depending on the FTD variant^70,71^. However, the proportion of right hemispheres sampled in this study was similar between controls (21%) and FTD (22%), suggesting that hemispheric differences likely did not significantly influence the results. Nonetheless, an investigation of different FTD variants that present either left- or right-dominant atrophy may be warranted to better understand this urea phenotype.

## 6 Conclusion

In conclusion, this study provides evidence of elevated urea levels in FTD and ALS *post-mortem* brain tissue, marking the 6th and 7th age-related dementia to exhibit this phenotype, and highlights urea as a potential universal therapeutic target for dementia. The elevated urea levels observed here and in other dementias may lead to increased carbamoylation and subsequent cell death. However, it remains unclear whether urea-induced carbamoylation could be a significant contributing factor to dementia pathogenesis or merely a byproduct of early pathogenic events. Future research should aim to elucidate the specific role of cerebral carbamoylation in the pathogenesis of dementia and identify the mechanisms underlying elevated brain urea levels.

## 7 Abbreviations

ALS: Amyotrophic lateral sclerosis
BBB: Blood-brain barrier
CNS: Central nervous system
FTD: Frontotemporal dementia
FTLD: Frontotemporal lobar degeneration
QC: Quality control
TDP-43: TAR DNA-binding protein of 43 kDa
UHPLC-MS/MS: Ultra-high performance-tandem mass spectrometry
UT-A: Urea transporter A
UT-B: Urea transporter B

## 8 Declarations

### 8.1 Ethics approval and consent to participate

Consent for the use of donated tissues for research and publication was collected by the NIH NeuroBioBank.

### 8.2 Consent for publication

Not applicable.

### 8.3 Data availability

All data generated or analysed during this study are included in this published article and its supplementary information files.

### 8.6 Authors’ contributions

SAP designed and performed experiments, analysed and interpreted data, and wrote the manuscript. SJC analysed and interpreted data. RDU interpreted the data and revised the manuscript. GJSC designed the experiments, interpreted the data, revised the manuscript, and bears overall responsibility for the integrity of this manuscript and of the study. All authors read and approved the final manuscript.

## 8.7 Acknowledgements

Human tissue was obtained from the NIH NeuroBioBank.

## Supplementary material

**Supplementary file 1**

.docx

Supplementary material

A single file containing all supplementary figures and tables.

**Supplementary file 2**

.xlsx

Raw brain urea values

A single file containing all raw brain urea values from both ALS and FTD case–control investigations.

