## Supplementary File 1 for "Elevated urea levels in human frontotemporal dementia and amyotrophic lateral sclerosis *post-mortem* brain tissue: Evidence of a multi-dementia pathogenic mechanism"

### Supplementary material

#### Supplementary Tables

**Supplementary Table 1. Selected reaction monitoring urea assay settings.**

| | Parent Ion Mass | Product Ion Mass | SRM Collision Energy | Retention Time (min) | S-Lens | Polarity |
|---|---|---|---|---|---|---|
| **Urea** | 61.1 | 44.1 | 18 | 1.10 | 34 | Positive |
| **Labelled Urea** | 63.1 | 45.1 | 18 | 1.10 | 34 | Positive |

**Supplementary Table 2. Individual patient characteristics for the Harvard Brain Tissue Resource Centre cohort.**

| BBN | Repository | Subject Sex | Hemisphere | Subject Age | PMI (hours) | Neuropathology Diagnosis | Genetic Diagnosis |
|---|---|---|---|---|---|---|---|
| S13898 | Harvard Brain Tissue Resource Centre | Female | Right | 72 | 18.25 | Diagnostic pathology not present | None Reported |
| S14415 | Harvard Brain Tissue Resource Centre | Female | Left | 68 | 21 | Diagnostic pathology not present | None Reported |
| S14826 | Harvard Brain Tissue Resource Centre | Male | Left | 75 | 20.5 | Diagnostic pathology not present | None Reported |
| **S14842*** | Harvard Brain Tissue Resource Centre | Female | Left | 71 | 20.3 | Diagnostic pathology not present | None Reported |
| S15795 | Harvard Brain Tissue Resource Centre | Female | Left | 74 | 18.5 | Diagnostic pathology not present | None Reported |
| S16461 | Harvard Brain Tissue Resource Centre | Female | Left | 69 | 23.33 | Diagnostic pathology not present | None Reported |
| S17567 | Harvard Brain Tissue Resource Centre | Female | Left | 66 | 23.17 | Diagnostic pathology not present | None Reported |
| S18190 | Harvard Brain Tissue Resource Centre | Female | Left | 47 | 25.83 | Diagnostic pathology not present | None Reported |
| S18670 | Harvard Brain Tissue Resource Centre | Female | Right | 73 | 26.92 | Diagnostic pathology not present | None Reported |
| S19397 | Harvard Brain Tissue Resource Centre | Female | Right | 60 | 28.15 | Diagnostic pathology not present | None Reported |
| S19871 | Harvard Brain Tissue Resource Centre | Female | Left | 66 | 23.2 | Diagnostic pathology not present | None Reported |
| S03755 | Harvard Brain Tissue Resource Centre | Male | Left | 76 | 18.17 | Diagnostic pathology not present | None Reported |
| S03893 | Harvard Brain Tissue Resource Centre | Male | Left | 66 | 24.55 | Diagnostic pathology not present | None Reported |
| S09190 | Harvard Brain Tissue Resource Centre | Female | Left | 71 | 20.5 | Diagnostic pathology not present | None Reported |
| S11641 | Harvard Brain Tissue Resource Centre | Female | Left | 75 | 24.5 | Diagnostic pathology not present | None Reported |
| S18399 | Harvard Brain Tissue Resource Centre | Female | Left | 78 | 25.21 | Frontotemporal dementia, unspecified | None Reported |
| S18571 | Harvard Brain Tissue Resource Centre | Female | Left | 64 | 25.88 | Frontotemporal dementia, unspecified | VCP / IBMPFD |
| S00288 | Harvard Brain Tissue Resource Centre | Female | Right | 77 | 15.83 | Frontotemporal dementia, unspecified | None Reported |
| S01450 | Harvard Brain Tissue Resource Centre | Male | Left | 67 | 25.25 | Frontotemporal dementia, unspecified | None Reported |
| **S02207**† | Harvard Brain Tissue Resource Centre | Female | Right | 84 | 19.25 | Frontotemporal dementia, unspecified | None Reported |
| S04746 | Harvard Brain Tissue Resource Centre | Female | Left | 62 | 27.75 | Frontotemporal dementia, unspecified | None Reported |
| S05494 | Harvard Brain Tissue Resource Centre | Male | Left | 61 | 19.41 | Frontotemporal dementia, unspecified | None Reported |
| S06963 | Harvard Brain Tissue Resource Centre | Female | Left | 80 | 26.17 | Frontotemporal dementia, unspecified | None Reported |
| S08631 | Harvard Brain Tissue Resource Centre | Female | Left | 57 | 23.72 | Frontotemporal dementia, unspecified | None Reported |
| | **Mean control** | 22% (M) | 21% (Right) | 70 | 22 | | |
| | **Mean FTD (PVC/FC cohort)** | 24% (M) | 22% (Right) | 70/68 | 23/24 | | |
| | **Brown–Forsythe and Welch ANOVA (PVC/FC cohort)** | | | >0.99/0.99 | 0.94/0.99 | | |

*One control was excluded due to suspected early-stage vascular dementia, which can independently alter urea concentrations. †One sample in the frontal cortex only was unable to be analysed due to an insufficient amount of tissue. Abbreviations: BBN: brain bank number; FC: frontal cortex; PMI: post-mortem interval; PVC: primary visual cortex.

**Supplementary Table 3. Individual patient characteristics for the University of Maryland Brain and Tissue Bank cohort.**

| BBN | Repository | Subject sex | Subject age | PMI (h) | Neuropathological diagnosis | Genetic diagnosis |
|---|---|---|---|---|---|---|
| 5604 | University of Maryland Brain and Tissue Bank | Female | 73 | 20 | Diagnostic pathology not present | None Reported |
| 6332 | University of Maryland Brain and Tissue Bank | Female | 95 | 10 | Diagnostic pathology not present | None Reported |
| 6357 | University of Maryland Brain and Tissue Bank | Male | 49 | 23 | Diagnostic pathology not present | None Reported |
| 6358 | University of Maryland Brain and Tissue Bank | Male | 37 | 26 | Diagnostic pathology not present | None Reported |
| 6365 | University of Maryland Brain and Tissue Bank | Male | 45 | 29 | Diagnostic pathology not present | None Reported |
| 6433 | University of Maryland Brain and Tissue Bank | Male | 47 | 22 | Diagnostic pathology not present | None Reported |
| 6574 | University of Maryland Brain and Tissue Bank | Male | 46 | 10 | Diagnostic pathology not present | None Reported |
| 6591 | University of Maryland Brain and Tissue Bank | Male | 35 | 22 | Diagnostic pathology not present | None Reported |
| 6592 | University of Maryland Brain and Tissue Bank | Female | 53 | 24 | Diagnostic pathology not present | None Reported |
| 6593 | University of Maryland Brain and Tissue Bank | Female | 48 | 26 | Diagnostic pathology not present | None Reported |
| **6688*** | University of Maryland Brain and Tissue Bank | Female | 106 | 15 | Diagnostic pathology not present | None Reported |
| 6753 | University of Maryland Brain and Tissue Bank | Male | 60 | 24 | Diagnostic pathology not present | None Reported |
| 6761 | University of Maryland Brain and Tissue Bank | Male | 56 | 22 | Diagnostic pathology not present | None Reported |
| **6819*** | University of Maryland Brain and Tissue Bank | Male | 80 | 22 | Diagnostic pathology not present | None Reported |
| 5782 | University of Maryland Brain and Tissue Bank | Female | 86 | 19 | Amyotrophic lateral sclerosis | None Reported |
| 5788 | University of Maryland Brain and Tissue Bank | Male | 83 | 22 | Amyotrophic lateral sclerosis | None Reported |
| **5789**† | University of Maryland Brain and Tissue Bank | Female | 71 | 17 | Amyotrophic lateral sclerosis | None Reported |
| 5858 | University of Maryland Brain and Tissue Bank | Female | 79 | 14 | Amyotrophic lateral sclerosis | None Reported |
| 5867 | University of Maryland Brain and Tissue Bank | Male | 51 | 19 | Amyotrophic lateral sclerosis | None Reported |
| 5868 | University of Maryland Brain and Tissue Bank | Male | 57 | 31 | Amyotrophic lateral sclerosis | None Reported |
| 5913 | University of Maryland Brain and Tissue Bank | Female | 60 | 25 | Amyotrophic lateral sclerosis | None Reported |
| 5974 | University of Maryland Brain and Tissue Bank | Male | 56 | 22 | Amyotrophic lateral sclerosis | None Reported |
| 6002 | University of Maryland Brain and Tissue Bank | Male | 77 | 14 | Amyotrophic lateral sclerosis | None Reported |
| 6349 | University of Maryland Brain and Tissue Bank | Male | 75 | 27 | Amyotrophic lateral sclerosis | None Reported |
| 6350 | University of Maryland Brain and Tissue Bank | Male | 60 | 10 | Amyotrophic lateral sclerosis | Amyotrophic lateral sclerosis |
| 6372 | University of Maryland Brain and Tissue Bank | Male | 45 | 8 | Amyotrophic lateral sclerosis | None Reported |
| 6377 | University of Maryland Brain and Tissue Bank | Female | 73 | 22 | Amyotrophic lateral sclerosis | None Reported |
| 6583 | University of Maryland Brain and Tissue Bank | Male | 66 | 19 | Amyotrophic lateral sclerosis | None Reported |
| | **Mean control** | 67% (M) | 54 | 22 | | |
| | **Mean ALS (DN/PMC cohort)** | 64/69% (M) | 67 | 19 | | |
| | **Brown–Forsythe and Welch ANOVA (DN/PMC cohort)** | | 0.17/0.20 | 0.91/0.95 | | |

*Samples were excluded in both dentate nucleus and primary motor cortex groups due to urea values being identified as outliers using box plots (Suppl. Fig?). †One sample was excluded in the primary motor cortex group only due the urea value being identified as an outlier. Abbreviations: BBN: brain bank number; DN: dentate nucleus; PMC: primary motor cortex; PMI: post-mortem interval.

**Suppl. Table 4. Sensitivity analyses for case–control urea investigations of FTD and ALS.**

| **FTD** | | |
|---|---|---|
| | **Frontal cortex** | **Primary visual cortex** |
| S-value (Based on Welch's T-test) | 11.5 | 6.6 |
| S-value (Based on Mann–Whitney U test) | 9.3 | 7.2 |
| E-value | 13.5 | 9.5 |
| Risk Ratio (RR) | 7.00 (1.94–25.30) | 5.00 (1.27–19.69) |
| RR CI E-value | 3.3 | 1.9 |
| Effect Size (Glass's delta) | 1.61 | 1.3 |
| **ALS** | | |
| | **Primary motor cortex** | **Dentate nucleus** |
| S-value (Based on Welch's T-test) | 4.6 | 3.4 |
| S-value (Based Mann–Whitney U test) | 6.0 | 2.2 |
| E-value | 10.6 | 5.4 |
| Risk Ratio (RR) | 5.54 (0.78–39.57) | 3.00 (0.76–11.80) |
| RR CI E-value | 1.0 | 1.0 |
| Effect Size (Glass's delta) | 1.1 | 1.7 |

**Suppl. Table 5. Multiregional sample size estimates for ALS and FTD case–controlled investigations.**

| **Brain region** | **Sample size estimate** |
|---|---|
| Frontal cortex (FTD/Con) | **12** |
| Primary visual cortex (FTD/Con) | 24 |
| Primary motor cortex (ALS/Con) | 72 |
| Dentate nucleus (ALS/Con) | 44 |

Required sample sizes were generated using a statistical power of 80% and an $\alpha$ error probability of 0.05. Values in bold represent groups that met the sample size criteria. Data were generated using G*Power (v. 3.1.9.4).

#### Supplementary Figures

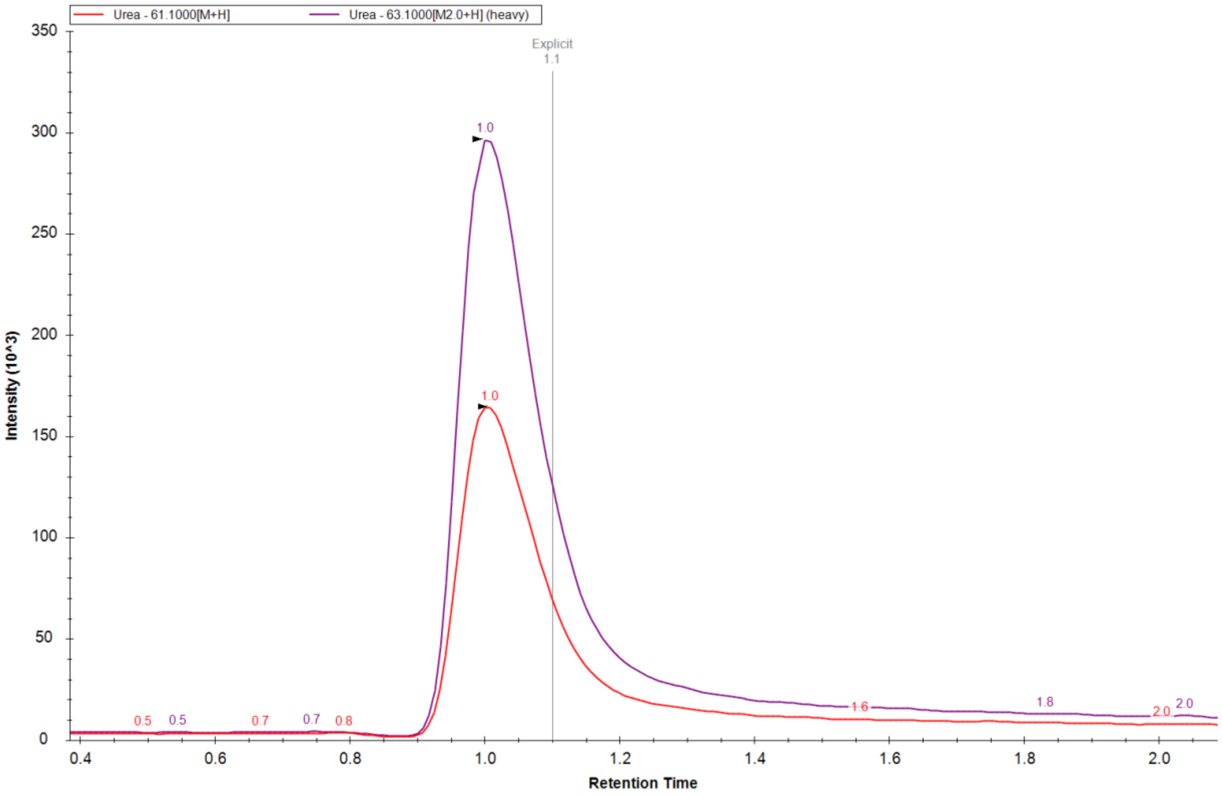


**Suppl. Fig. 1.** A representative chromatogram of the selected reaction monitoring urea assay in human brain tissue. The red peak represents endogenous urea, and the purple peak represents heavy-labelled urea.

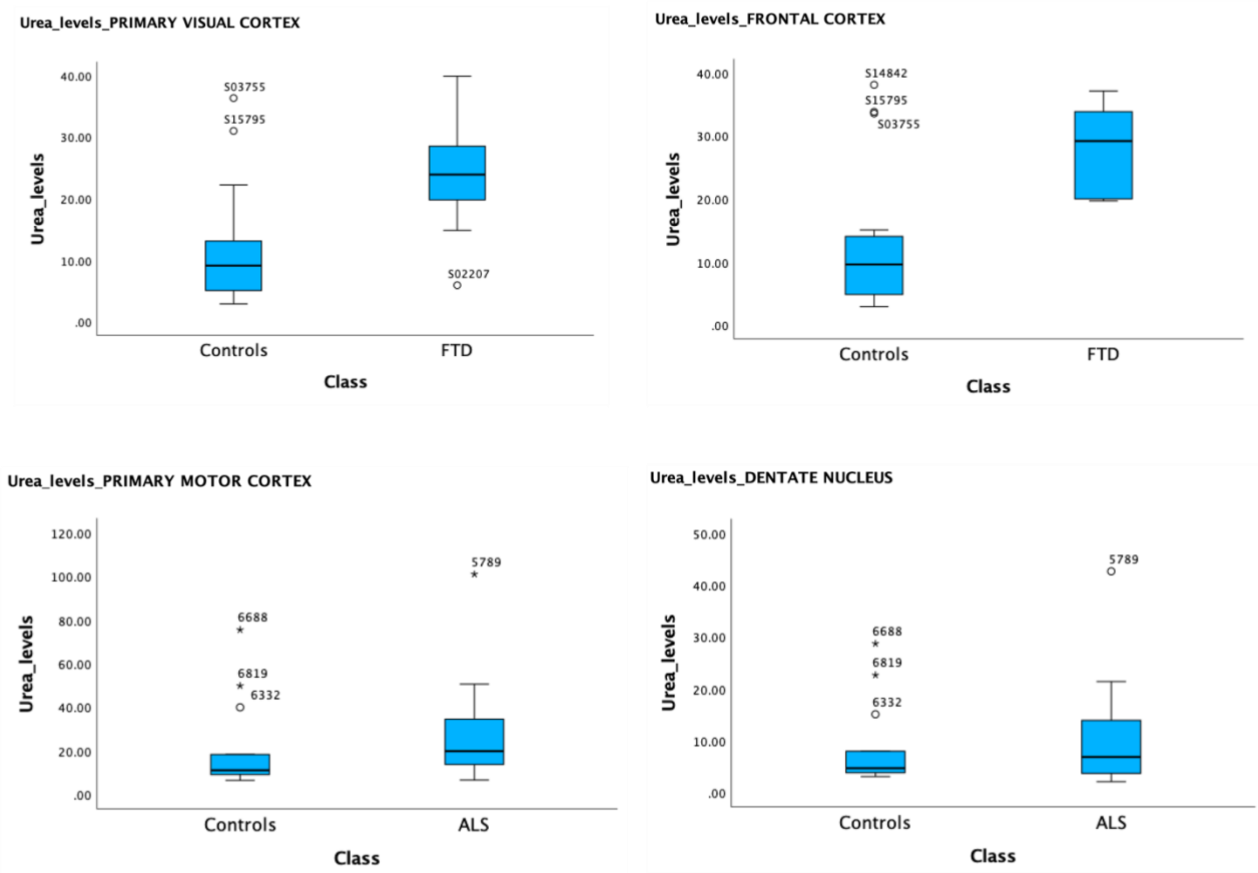


**Suppl. Fig. 2.** *Post-mortem* brain urea levels from FTD and ALS case–control investigations fitted in boxplots to identify possible outliers. Extreme outliers were defined as being more than 3 interquartile range (IQR) below the first quartile or above the third quartile (represented as an asterisk). Mild outliers were defined as being more than 1.5 IQR below the first quartile or above the third quartile (represented as a circle). Boxplots were generated using SPSS (version 29.0.0.0).

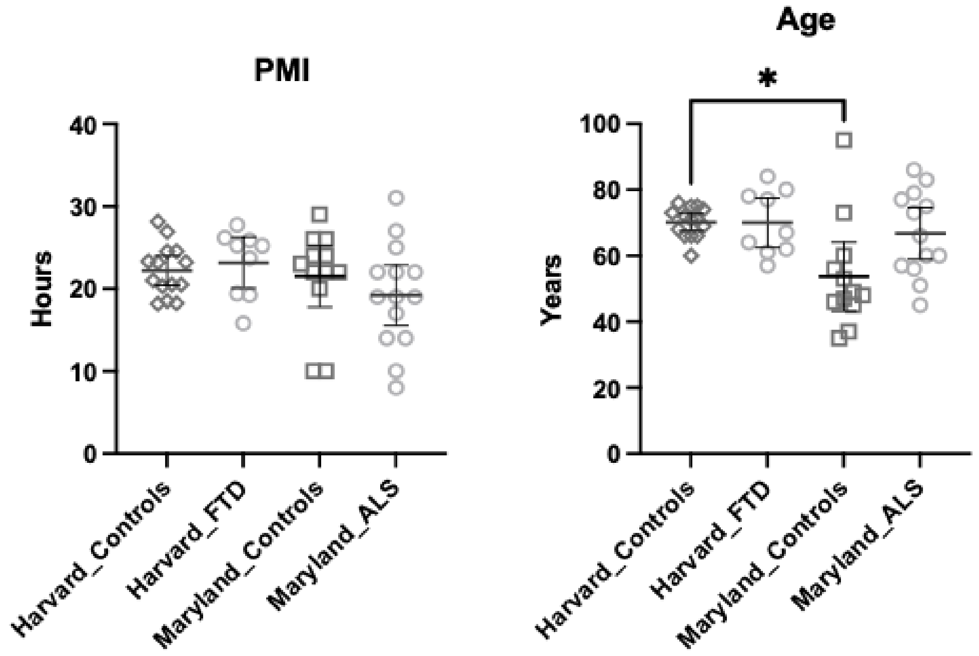


**Suppl. Fig. 3.** Cohort variable (PMI and age) comparisons across all tissue groups (means ±95% CI). Cohort variable differences were determined using Brown–Forsythe and Welch ANOVA with Dunnett's T3 multiple comparisons test. **p* $< 0.0332$.

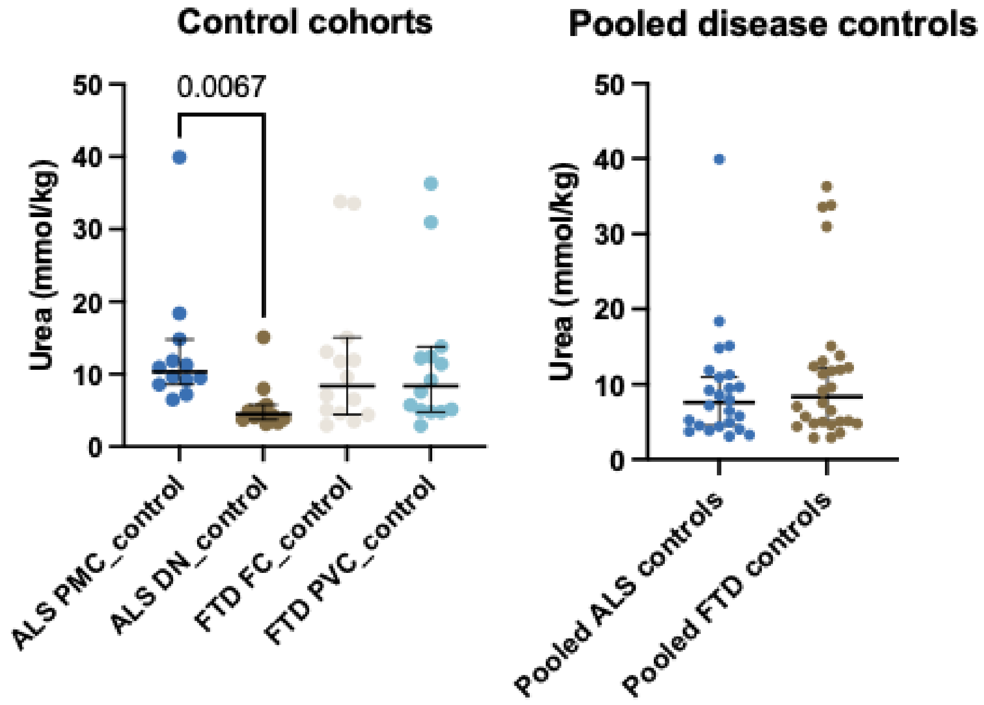


**Suppl. Fig. 4.** Comparison of urea levels between control cohorts. Cohort differences were determined using a Kruskal–Wallis test with Dunn's multiple comparison test or a Mann–Whitney U test.

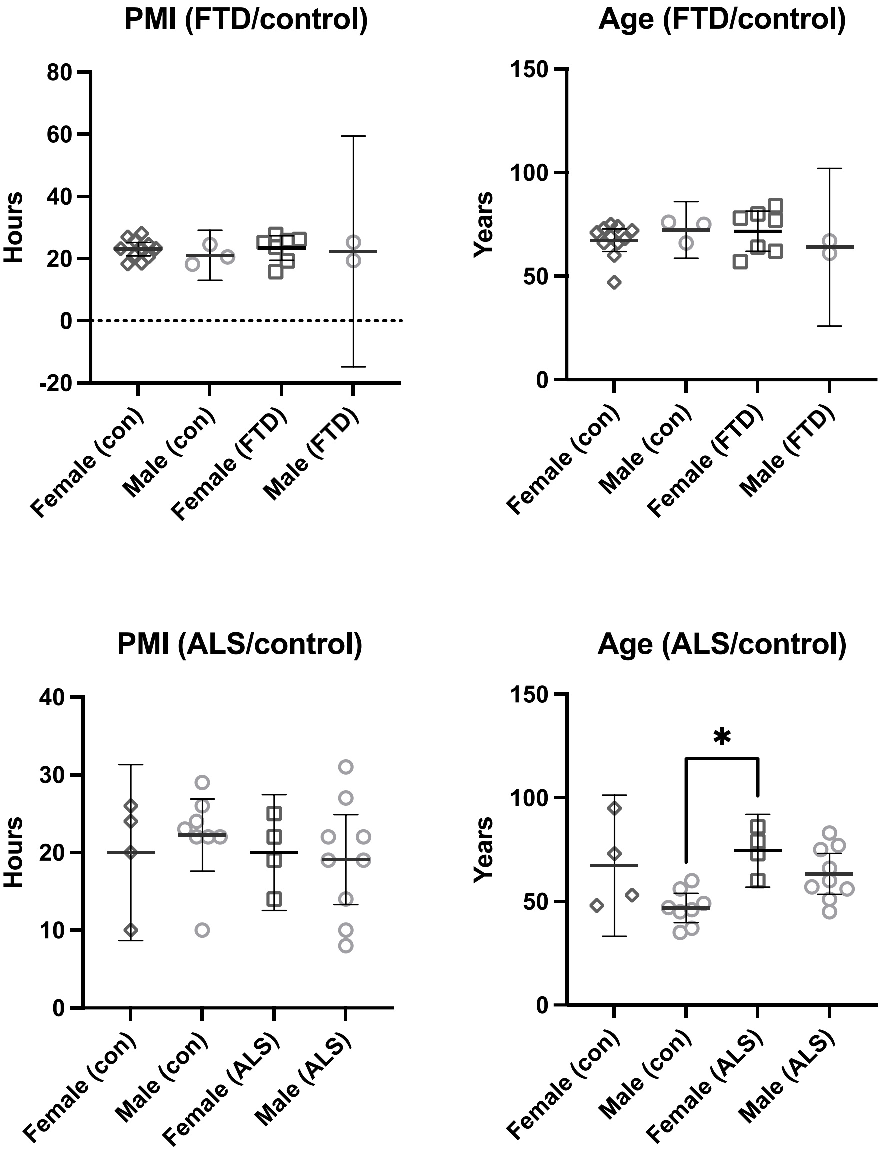


**Suppl. Fig. 4.** Cohort variable (PMI and age) comparisons (means ±95% CI) separated by sex for FTD and ALS case–control investigations. Cohort variable differences were determined using the Kruskal–Wallis test with Dunn's multiple comparisons test. *$p < 0.0332$.

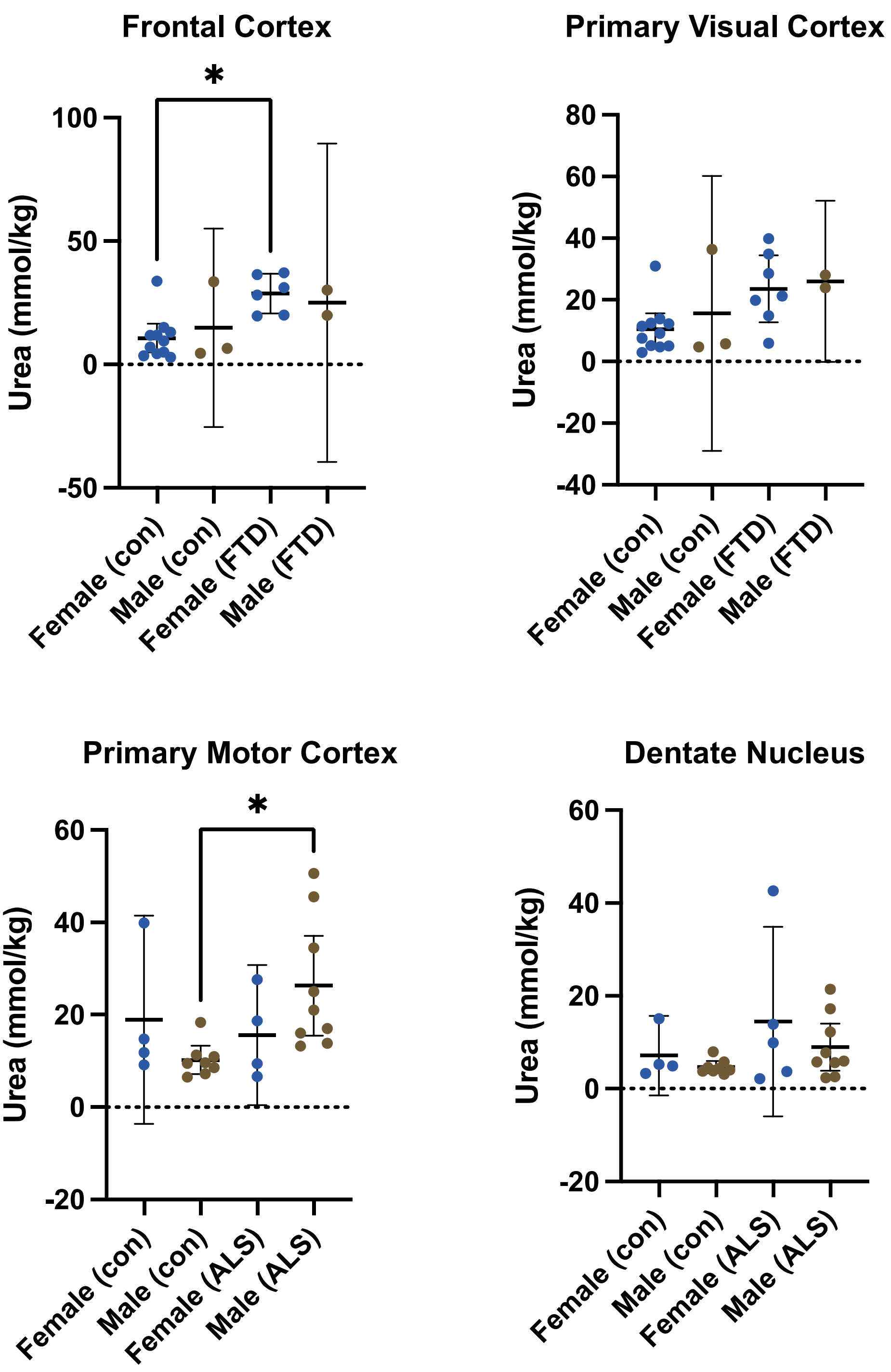


**Suppl. Fig. 5.** Regional brain urea concentrations separated by sex (means ±95% CI). Brain urea differences were determined using the Kruskal–Wallis test with Dunn's multiple comparisons test. *$p < 0.0332$.
